# Estimation of universal and taxon-specific parameters of prokaryotic genome evolution

**DOI:** 10.1101/137430

**Authors:** Itamar Sela, Yuri I. Wolf, Eugene V. Koonin

## Abstract

Our recent study on mathematical modeling of microbial genome evolution indicated that, on average, genomes of bacteria and archaea evolve in the regime of mutation-selection balance defined by positive selection coefficients associated with gene acquisition that is counter-acted by the intrinsic deletion bias. This analysis was based on the strong assumption that parameters of genome evolution are universal across the diversity of bacteria and archaea, and yielded extremely low values of the selection coefficient. Here we further refine the modeling approach by taking into account evolutionary factors specific for individual groups of microbes using two independent fitting strategies, an ad hoc hard fitting scheme and an hierarchical Bayesian model. The resulting estimate of the mean selection coefficient of *s*∼10^-10^ associated with the gain of one gene implies that, on average, acquisition of a gene is beneficial, and that microbial genomes typically evolve under a weak selection regime that might transition to strong selection in highly abundant organisms with large effective population sizes. The apparent selective pressure towards larger genomes is balanced by the deletion bias, which is estimated to be consistently greater than unity for all analyzed groups of microbes. The estimated values of *s* are more realistic than the lower values obtained previously, indicating that global and group-specific evolutionary factors synergistically affect microbial genome evolution that seems to be driven primarily by adaptation to existence in diverse niches.

## Introduction

Prokaryotes have compact genomes, in terms of the number of genes and especially gene density, with typically short intergenic regions comprising less than 10% of the genome (Koonin and Wolf, 2008; Lynch and Conery, 2003; Mira et al., 2001). Deciphering the evolutionary forces that keep prokaryotic genome compact is an important problem in evolutionary biology. The common view, steeped in a population-genetic argument, seems to be that selection favors compact genomes in the fast-reproducing prokaryotes with large effective population sizes, to minimize the replication time and the energetic burden that is associated with gene expression (Lynch and Conery, 2003; Lynch and Marinov, 2015). This theory provides a plausible explanation for the observed dramatic differences in the typical size and architecture between prokaryotic and eukaryotic genomes, with the latter being up to several orders of magnitude larger than the former and, in many case, containing extensive non-coding regions (Koonin, 2009). Under the population-genetic perspective, the large effective population sizes of prokaryotes enhance the selection pressure and allow efficient elimination of superfluous genetic material (Lynch, 2007, 2006; Lynch and Conery, 2003; Lynch and Marinov, 2015).

The population-genetic theory predicts an inverse correlation between genome size and the strength of selection, and this prediction generally holds across the full range of genome sizes, from viruses to multicellular eukaryotes (Lynch, 2007; Lynch and Conery, 2003). However, a detailed analysis of the relationship between the genome size and selection strength within prokaryotes reveals the opposite trend: genome size correlates positively and significantly with the protein-level selection strength indicating that larger genomes are typically subject to stronger selection on the protein level (Kuo et al., 2009; Novichkov et al., 2009b; Sela et al., 2016). The protein-level selection is measured by the ratio of non-synonymous to synonymous mutation rates (d*N*/d*S* ratio) (Hurst, 2002) in core genes that are common across (nearly) all prokaryotes (Koonin, 2003). The underlying assumption is that the effects of single non-synonymous mutations in these core, functionally conserved genes are similar (associated with similar selection coefficients) across all prokaryotes (Sela et al., 2016). The differences in the observed d*N*/d*S* values between groups of prokaryotes are accordingly assumed to reflect differences in selection strength. At least formally, within the population-genetic theory, this assumption translates to similar selection coefficients but different effective population sizes.

Recently, we performed an analysis of the factors that govern prokaryotic genome size evolution by developing a population-genetic evolutionary model and testing its predictions against the genome size distributions in 60 groups of closely related bacterial and archaeal genomes (Sela, Wolf et al. 2016). Within the modeling framework, the genome size evolution is represented as stochastic gain and loss of genes, an approach that is motivated by the dominant role of horizontal gene transfer in microbial evolution (Doolittle, 1999; Koonin et al., 2001; Pal et al., 2005; Puigbo et al., 2014; Treangen and Rocha, 2011). Specifically, the model predicts a distribution of the genome sizes for the given values of the effective population size, the deletion bias and the selection coefficient associated with the gain of a gene. Using maximum-likelihood optimization methods, the values of the deletion bias and the selection coefficients can be inferred from the data. Under the simplifying assumption that the mean selection coefficients and deletion bias are similar across the diversity of prokaryotes, the global mean values of these factors can be used in the model. Under this assumption, the different observed mean genome sizes among prokaryotic groups are due to the differences in the effective population sizes (*N*_*e*_). The model then predicts a global trend line, which represents the dependency of the mean genome size on the effective population size. More realistically, however, the selection coefficients and the deletion bias values can differ between prokaryotic groups, and the observed genome sizes therefore deviate from the global trend. The challenge is to account for such deviations as fully as possible, without discounting the effect of the universal behavior.

In our previous study (Sela et al., 2016), the fitting of the data to the model was performed in two stages: first, global parameters were fitted, and at the second stage, some parameters were taken as latent variables and were optimized to maximize the log-likelihood. This methodology is most accurate when deviations from the global trend are small compared to the distribution width. Here, we substantially modify the fitting procedure, to account for the specific factors affecting the genome evolution in different groups of prokaryotes, without obscuring the global trend. The resulting parameters of microbial evolution appear to be more realistic than those obtained with the previous, simplified approach.

## Material and Methods

### Genomic dataset and estimation of selection pressure and effective population size

A dataset of 707 bacterial and archaeal genomes clustered in 60 groups of closely related organisms was constructed using the Alignable Tight Genomic Cluster (ATGC) database (Kristensen et al., 2017; Novichkov et al., 2009a). For simplicity, these individual genomes will be referred to as “species” although many of them represent strains and isolates within the formally described microbial species. In addition to the genome size, which is known for all species in the database, a characteristic value of selection strength was assigned to each cluster (see Figure 1A). Selection strength was inferred on the protein level, by estimating the d*N*/d*S* ratio of 54 core gene families that are common for all or nearly all prokaryotes. Specifically, these alignments of the core proteins constructed using the MUSCLE program (Edgar, 2004) were concatenated, converted to the underlying nucleotide sequence alignments, and the characteristic d*N*/d*S* value (Yang, 2007) for each cluster was estimated as the median d*N*/d*S* for all species pairs in the cluster. As shown previously, the median *dN/dS* is a stable characteristic of an ATGC that is robust to variations in the set of genome pairs employed for the estimation (Novichkov et al., 2009b). The effective population size *N*_*e*_ for each cluster is deduced from the typical d*N*/d*S* value, using the approach developed by Kryazhimskiy and Plotkin (Kryazhimskiy and Plotkin, 2008). The effective population size calculation is performed under the following assumptions. Core genes are assumed to evolve under the weak mutation limit regime, where the mutation rate is low such that mutations appear sequentially. In addition, it is assumed that synonymous mutations are strictly neutral, and that the selection coefficient associated with non-synonymous mutations is similar for all core genes in all prokaryotes. Finally, the selection coefficient value of non-synonymous mutations is set such that the effective population size for ATGC001, that contains *Escherichia coli* strains, is 10^9^ and the effective population size for all other clusters is calculated accordingly.

**Figure 1.**
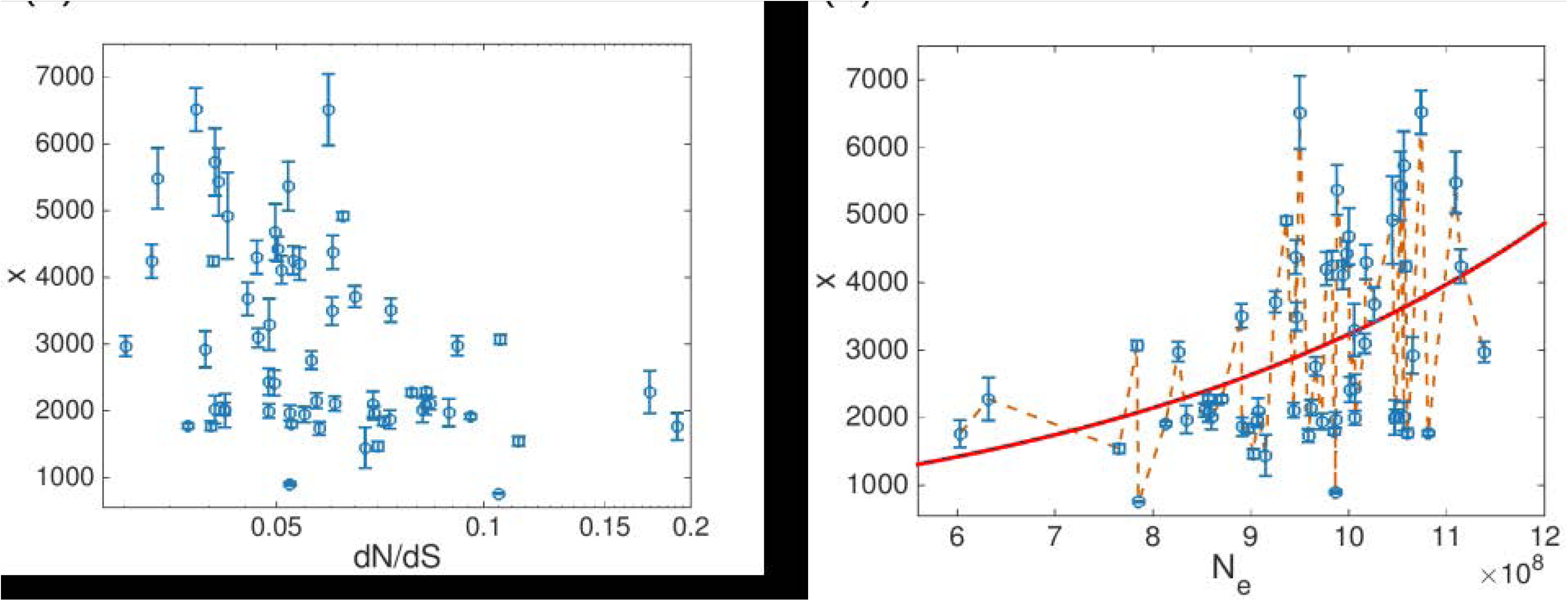
Genome size and selection strength in prokaryotes. **(A)** Mean number of genes *x* is plotted against inferred selection strength d*N*/d*S* where each point represent one prokaryotic cluster (ATGC). Error bars represent genome sizes distributions widths and indicate one standard deviation. **(B)** Mean number of genes is plotted against extracted effective population size *N*_*e*_. A representative global trend line of mean genome size as predicted by the model (see Eq.(8)), where all model parameters are assumed to be global ***θ*** = {*s*, *r*′, *λ*} is indicated by a red line. The approach implemented in the hard fitting methodology, where Eq.(8) is used in order to set latent variable value such that model distributions are centered around observed genome sizes, is illustrated in a dashed orange line.

### Maximum-likelihood framework for model parameters optimization

The objective is to infer the unknown parameters of the genome size model presented below from the genomic dataset. The probability of a set of observations ***X***, namely, observed genome sizes in all species in all ATGCs, is given by a distribution predicted by the genome size population model. The distribution depends on two types of parameters: known parameters ***Z***, and unknown parameters ***θ***. For the genome size population model, the known parameter is the effective population size *N*_*e*_, which is calculated for each ATGC. The unknown parameters are deletion bias (*r)* and selection coefficient (*s*) associated with the gain of a single gene. Simply put, the goal is to optimize ***θ*** by fitting the model distribution to the observed genome sizes in terms of log-likelihood. Optimization is performed by maximizing ℓ(***θ***)

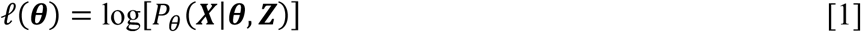

The calculation of *P*_*θ*_(***X***|***θ***, ***Z***) from the genome size population model is presented in detail in the Results section.

## Results

### Global model of genome evolution

The mean genome sizes and the d*N*/d*S* values correlate negatively and significantly, with Spearman’s rank correlation coefficient *ρ* = −0.397 and *p*-value 0.0017, in agreement with the previous observations (Kuo et al., 2009; Novichkov et al., 2009b; Sela et al., 2016)(Figure 1A). Effective population sizes are extracted from the d*N*/d*S* values for each ATGC, resulting in the same correlation, but with the opposite sign, between genome size *x* and *N*_*e*_. These correlations indicate that the genome size is determined, to a large extent, by global evolutionary factors that are shared by all prokaryotes. On top of the global factors, there obviously are local influences, such as different lifestyles, environments and availability of genetic material. The goal of the present work is to accurately assess the global factors that govern genome size evolution and are partially masked by local effects, and additionally, to compare the local factors for different groups of bacteria and archaea.

Evolution of prokaryotic genomes can be described within the framework of population genetics by a stochastic process of gene gain and loss events (Sela et al., 2016). In brief, a genome is modeled as a collection of *x* genes, where genome size is assumed to evolve through elementary events of acquisition or deletion of one gene at a time, occurring with rates *α* and *β*, respectively. Genes are assumed to be acquired from an infinite gene pool. Gene gains and losses are either fixed or eliminated stochastically, with a fixation probability *F*. In the weak mutation limit, the fixation probability can be expressed as (McCandlish et al., 2015)

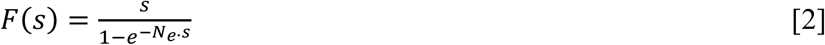

where *N*_*e*_ is the effective population size and *s* is the selection coefficient associated with acquisition of a single gene. That is, assuming that the reproduction rate for genome of size *x* is 1, the reproduction rate for a genome of size *x* + 1 is 1 + *s*. To obtain the selection coefficient associated with deletion of a gene, the event of gene deletion is considered: the reproduction rate for genome size *x* + 1 is set as 1, and the reproduction rate for genome size *x* can be therefore approximated by 1 - *s*, so that

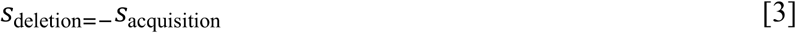

The gain rate, *P*_+_, is given by the multiplication of the acquisition rate *α*, and the fixation probability of a gene acquisition event. In general, both the acquisition rate and the selection coefficient associated with the acquisition of a gene depend on the genome size:

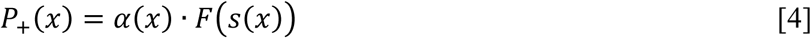

Using the relation *s*_deletion=−_*s*_acquisition_ derived above, we get a similar expression for the loss rate, denoted by *P*_−_

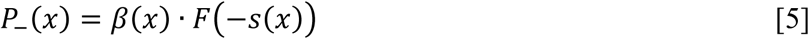

Genome size dynamics is then a chain of stochastic gain and loss events, and can be described by the equation

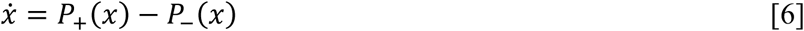

If for a some value of *x*, denoted *x*_0_, gain and loss rates are equal, i.e. the evolving genome fluctuates stochastically around this value (under a condition discussed below, see Eq.(10)), the dynamics of Eq.(6) implies a steady state distribution *f*(*x*) of the genomes sizes. This distribution has an extremum at *x*_0_, and is given by

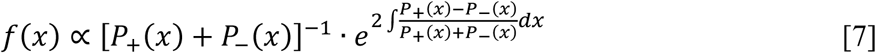

If the distribution is symmetric, *x*_0_ is the mean genome size, and given that *f*(*x*) is only slightly skewed with relevant model parameters (see Figure 2), *x*_0_ is taken as an approximation for the mean genome size. With respect to the model parameters, *x*_0_ satisfies the relation

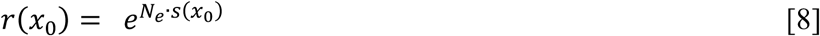

where *r*(*x*) is the deletion bias, defined as the ratio of the deletion and acquisition rates:

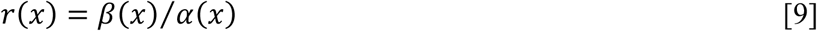

**Figure 2.**
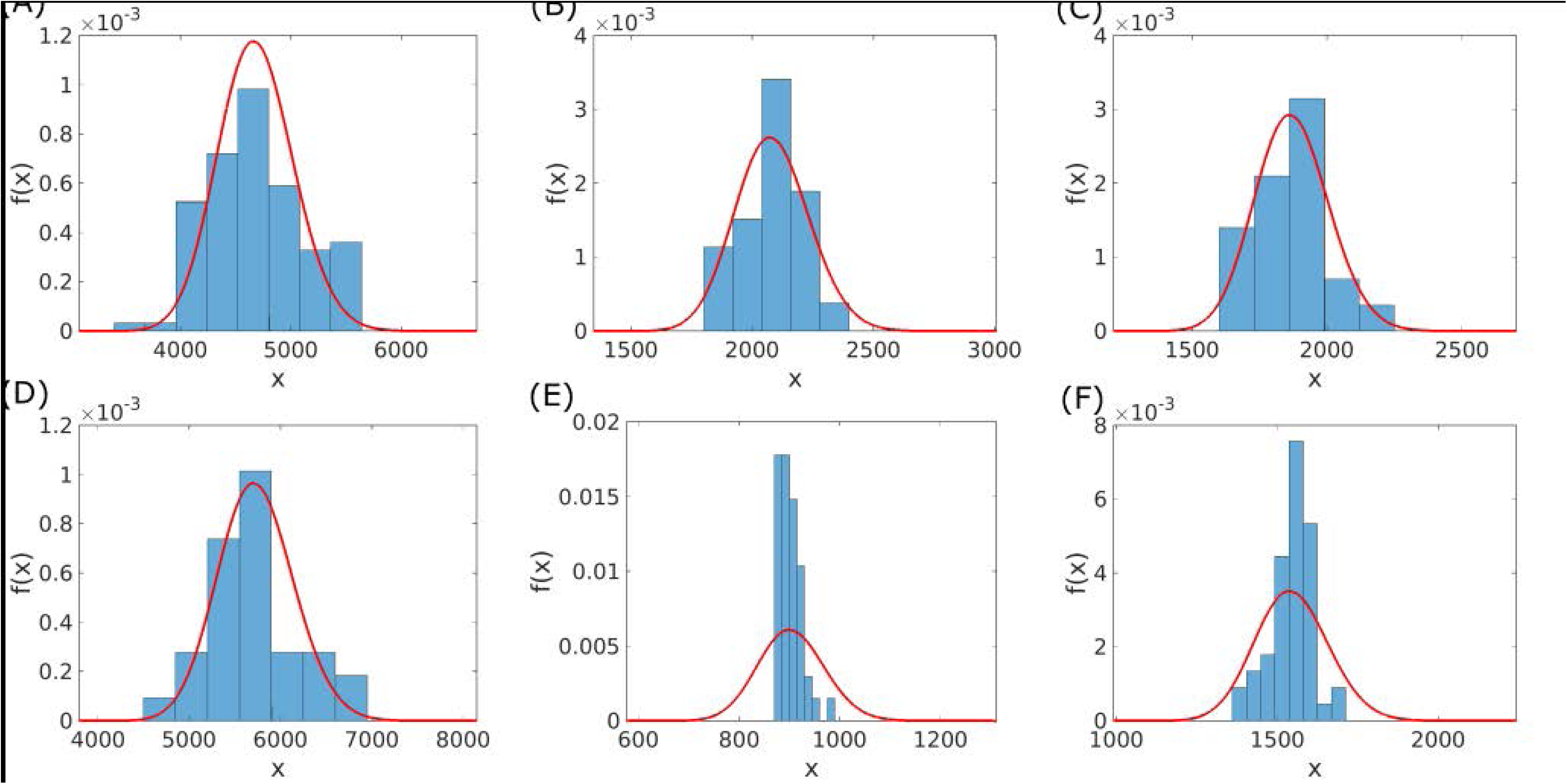
Comparison of the observed and model-generated genome size distributions for 6 ATGCs that consist of at least 20 species. Empirical genome sizes are indicated by bars and model distributions by red solid lines. For model distributions Eq.(7) was used, together with the deletion bias of Eq.(17). Model parameters were optimized using the hierarchical Bayesian model method, with the linear coefficient *a* of the acquisition rate (see Eq.(15)) as latent variable. Optimized parameters are listed in Table 2 and in Supplementary table 2. The ATGCs are as follows (the numbers of genomes for each ATGC are indicated in parentheses): (A) ATGC0001 (109), (B) ATGC0003 (22), (C) ATGC0004 (22), (D) ATGC0014 (31). (E) ATGC0021 (45) and (F) ATGC0050 (51)

The extremum point of *f*(*x*) at *x*_0_ can be either a maximum or a minimum. The case where *f*(*x*) has a minimum at *x*_0_ corresponds to genomes that are either collapsing or growing infinitely, and is biologically irrelevant. The extremum point at *x*_0_ is a maximum when

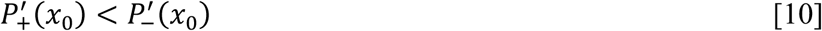

**Table 2.** Optimal fits for the genome evolution model parameters using the linear model of deletion bias (Eq.(17))

| Methodology | $\varphi$ | $s$ | $a$ | $b$ | $\ell(\theta)$ | $R^2$ | KS $p$-value | $\varphi_0$ | $\sigma_\varphi$ | $\rho$ | $\rho$<br>$p$-value |
| --- | --- | --- | --- | --- | --- | --- | --- | --- | --- | --- | --- |
| H | $s$ | - | 0.810 | 186 | -4700 | 0.175 | 0.52 | $1.26 \cdot 10^{-10}$ | $2.8 \cdot 10^{-11}$ | -0.01 | 0.92 |
| B | | | 0.825 | 167 | -4913 | - | - | $1.18 \cdot 10^{-10}$ | $2.5 \cdot 10^{-11}$ | -0.04 | 0.79 |
| H | $a$ | $1.41 \cdot 10^{-10}$ | - | 187 | -4696 | 0.175 | 0.4 | 0.80 | 0.04 | -0.02 | 0.88 |
| B | | $1.28 \cdot 10^{-10}$ | | 167 | -4909 | - | - | 0.816 | 0.03 | -0.03 | 0.79 |
| H | $b$ | $1.30 \cdot 10^{-10}$ | 0.824 | - | -4759 | 0.175 | 0.35 | 174 | 77 | 0.01 | 0.92 |
| B | | $1.91 \cdot 10^{-10}$ | 0.782 | | -4944 | - | - | 148 | 68 | -0.24 | 0.06 |

**H**, hard fitting methodology; **B**, hierarchical Bayesian model fitting.

Finally, explicit functional forms for *s*(*x*), *α*(*x*) and *β*(*x*) are assumed in the fitting process. The selection coefficient is taken as constant with respect to genome size

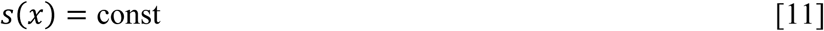

and two forms of acquisition and deletion rates are considered. The first corresponds to the deletion bias in the form of a power law

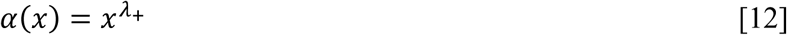

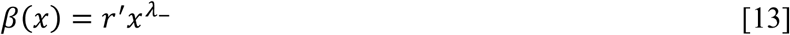

with

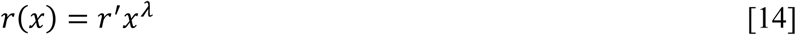

where *λ* = *λ*_−_ − *λ*_+_; because the distribution given by Eq.(7) is not sensitive to *λ*_+_ values, it was set to the value of 10^-3^. In addition, a linear model was considered, where

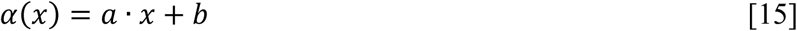

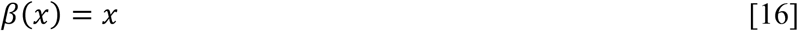

and the deletion bias is then given by

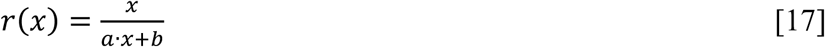

The selection coefficient was taken as constant (independent of genome size) for simplicity. Preliminary calculations with additional linear term in genome size gave similar results, both in terms of the log likelihood and fitted parameter values (see Table S1) The deletion bias is modelled by a power law with respect to genome size because it encompasses the two extreme cases of constant or linear dependence, along with all intermediates. For compatibility with birth-death-transfer models, in which linear acquisition and deletion rates are assumed (Iranzo et al., 2017), the deletion bias given by Eq.(17) was studied as well. With the formulations given above, the population model for genome size evolution contains one known parameter, *N*_*e*_, and a set of three unknown parameters: either {*s*, *r*′, *λ*} or {*s*, *a*, *b*}, depending to the choice of the model for the acquisition and deletion rates.

### Group-specific factors in prokaryotic genome evolution

The assumption that all model parameters are universal across the diversity of prokaryotes translates into a global trend line (see Figure 1B) because in this case, groups of prokaryotic species differ from each other only by the typical effective population size. However, when the model parameters are fitted under the assumption that all unknown parameters are universal, the observed distributions of the microbial genome sizes are much wider than the distributions predicted by the model (see Figure 3A) indicating that ATGC-specific factors play a non-negligible role in genome evolution. Deviations from the global trend line due to local effects can be incorporated into the model by introducing a latent variable ***φ***, i. e. assigning ATGC-specific values to one of the model parameters. The underlying assumption is that the universal dependency of the genome size on the effective population size is captured by the global parameters ***θ***, whereas the deviations from the universal behavior caused by ATGC-specific effects are incorporated in the model through different values of a latent variable ***φ***. Because variation in one parameter of the model can be compensated by variation in a different parameter (e.g. the *s* value can be adjusted to compensate for variation in *r*′ resulting in the same distribution; see Figure S2), standard methods for latent parameters fitting are not applicable.

**Figure 3.**
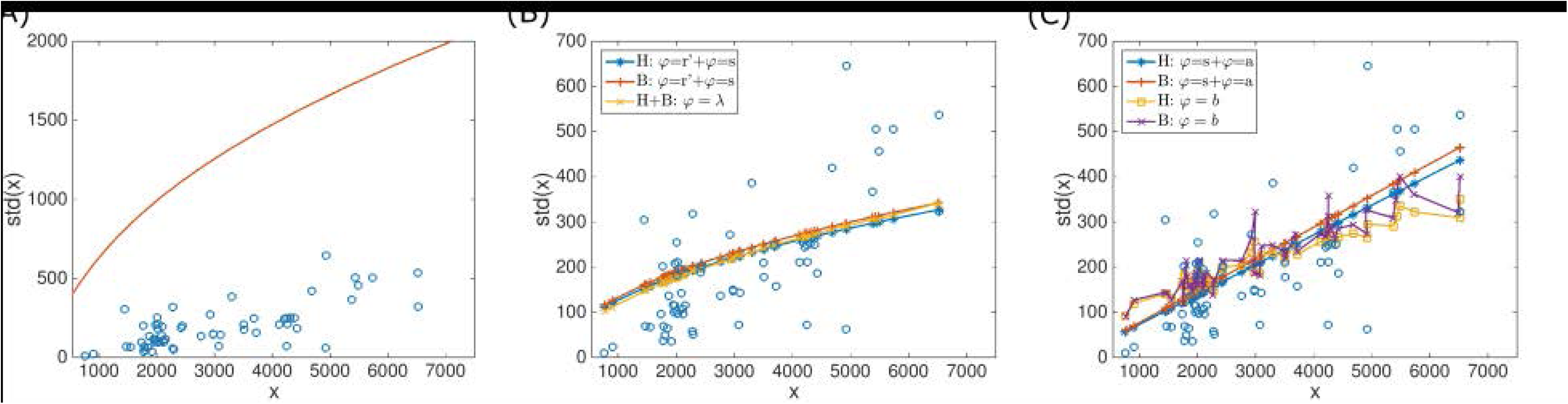
Prokaryotic genome size distribution width plotted vs. genome size. The standard deviation is taken as the proxy for the distribution width. ATGCs are indicated by circles and model fits by lines. (A) Model prediction using the deletion bias of Eq.(14) with parameters optimized under the assumption that all three model parameters as universal (Sela et al., 2016). (B) Six model fits with the deletion bias of Eq.(14) (fitted parameters are given in Table 1). In all fits, one model parameter was set as a latent variable. The model parameter that was set as a latent variable and the methodology used for fitting are indicated in the inset; fits that were visually indistinguishable are represented by the same line. H, hard fitting method; B, hierarchical Bayesian model. (C) Same as panel B, for the deletion bias of Eq.(17) (fitted parameters are given in Table 2).

Therefore, we developed two fitting methodologies: i) an *ad hoc* hard-fitting algorithm and ii) an hierarchical Bayesian fitting procedure. In both methodologies, ATGC-specific ***φ*** values are assigned according to the ***θ*** values. The probability of the observed genome sizes, *P*_*o*_(***X***|***θ***, ***φ***, ***Z***), is calculated numerically using the steady state genome size distribution *f*(*x*) of Eq.(7), as explained below.

**Table 1.** Optimal fits for the genome evolution model parameters using the power law model of deletion bias (Eq.(14))

| Methodology | $\varphi$ | $s$ | $r'$ | $\lambda$ | $\ell(\theta)$ | $R^2$ | KS $p$-value | $\varphi_0$ | $\sigma_\varphi$ | $\rho$ | $\rho$<br>$p$-value |
| --- | --- | --- | --- | --- | --- | --- | --- | --- | --- | --- | --- |
| H | $s$ | - | 0.693 | 0.061 | -4782 | 0.179 | 0.35 | $1.20 \cdot 10^{-10}$ | $2.8 \cdot 10^{-11}$ | -0.06 | 0.67 |
| B | | | 0.703 | 0.056 | -4975 | - | - | $9.0 \cdot 10^{-11}$ | $2.5 \cdot 10^{-11}$ | 0.04 | 0.78 |
| H | $r'$ | $1.25 \cdot 10^{-10}$ | - | 0.061 | -4782 | 0.179 | 0.35 | 0.70 | 0.018 | 0.03 | 0.83 |
| B | | $1.01 \cdot 10^{-10}$ | | 0.056 | -4975 | - | - | 0.710 | 0.017 | -0.02 | 0.87 |
| H | $\lambda$ | $1.27 \cdot 10^{-10}$ | 0.688 | - | -4770 | 0.179 | 0.32 | 0.0628 | 0.004 | 0.03 | 0.80 |
| B | | $8.7 \cdot 10^{-11}$ | 0.666 | | -4924 | - | - | 0.062 | 0.003 | -0.1 | 0.42 |

**H**, hard fitting methodology; **B**, hierarchical Bayesian model fitting.

The distributions produced by the model under optimized parameters are compared to the observed distributions in Figure 2. First, latent variable values are set for each ATGC, such that values are assigned to all three unknown model parameters. The details of this stage are discussed below. For each ATGC, acquisition and deletion rates are then calculated, using either Eqs.(12) and (13), or Eqs.(15) and (16). Together with the fixation probability, which is given by Eq.(2) and calculated using the ***θ*** and *Z* values, the acquisition and deletion rates are used to calculate the gain and loss rates of Eqs.(4) and (5). The gain and loss rates are then substituted into Eq.(7), and the genome size distribution is calculated and normalized numerically. Finally, the probability of the observed genome sizes is given by the product of the distribution values at the observed genome sizes ***X***

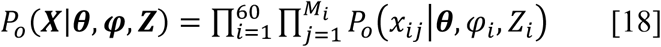

where *x*_*ij*_ is observed genome size for species *j* out of *M*_*i*_ species of ATGC *i*, and *φ*_*i*_ and *Z*_*i*_ are ATGC-specific values of the latent variable and effective population size, respectively. For example, when setting the linear coefficient *a* of the acquisition rate of Eq.(15) as the latent variable, we have

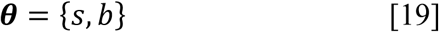

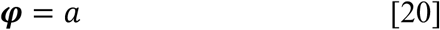

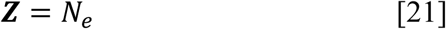

For given *s* and *b* values, an ATGC-specific value is assigned for *a*, such that values are assigned to all model parameters and *P*_*o*_(***X***|***θ***, ***φ***, ***Z***) can be calculated following the steps described above.

In the *ad-hoc* fitting procedure, one model parameter is set as a latent variable, and the two remaining unknown model parameters are considered global and included in ***θ***. Eq.(8) is used to adjust the latent variable value according to the ***θ*** values and center model distributions around data points (see Figure 1B)

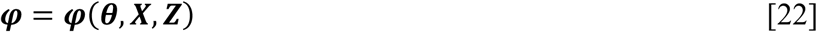

The log-likelihood is then calculated using Eq.(1) with

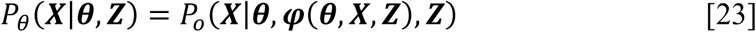

and *P*_o_(***X***|***θ***, ***φ***(***θ***, ***X***, ***Z***), ***Z***) is calculated using Eq.(7) as explained above. However, different values of global parameters ***θ*** can be compensated by the value of the latent variable ***φ*** to yield similar genome size distributions (see Figure S2). Therefore, an additional constraint is applied to the ***θ*** values in the optimization procedure and combined with the log likelihood ℓ(***θ***) of Eq.(1). The global parameters ***θ*** represent the universal evolutionary factors that entail the observed genome size and effective population size correlation. It is therefore natural to use in the optimization not only the log-likelihood but also the goodness of fit of the global trend line associated with the ***θ*** values. The global trend is produced using Eq.(8) by assuming that all three model parameters are universal; however, under this optimization methodology, ***θ*** is a set of only two global model parameters. The set of global parameters ***θ*** is therefore completed by a single representative value of the latent variable, denoted ⟨*φ*⟩, to produce the global trend line. The goodness of fit is then given by the *R*^2^ value for the global trend line and mean genome sizes of the different ATGCs (see Figure 1B). The *R*^2^ value clearly depends not only on the values of the two universal model parameters ***θ***, but also on the value of ⟨*φ*⟩. For the optimization of ***θ*** values, the maximum possible *R*^2^ value for the given ***θ*** values is taken.

The goodness of fit for the global trend line is optimized together with the log likelihood, by minimizing a goal function *G*(***θ***):

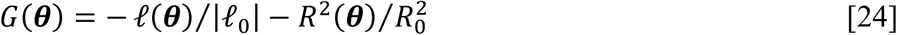

where the log-likelihood and goodness of fit are normalized to give comparable values. Specifically, the values |ℓ_0_| = 4773 and 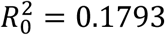 were used as these are close to the optimal values of log-likelihood and goodness of fit, respectively, for our data set. Fitting was performed for all three assignments of the latent parameter and the two representations of the deletion bias, namely, *φ* = *s*, *φ* = *λ* and *φ* = *r*′ for the deletion bias of Eq.(14), and *φ* = *s*, *φ* = *a* and *φ* = *b* for the deletion bias of Eq.(17). In all 6 cases, the results were similar, in terms of both the optimized values of the selection coefficient and log-likelihood. The results are summarized in Tables 1 and 2, and the fitted latent variable values are shown in Figures 4 and 5. Notably, there was no significant correlation of the fitted latent variable values and effective population size (Tables 1 and 2), suggesting that the universal correlation between the genome size and the effective population size is not masked by assigning ATGC-specific value to model parameters using this approach. For comparison with the hierarchical Bayesian model approach (see below), the optimized latent variable values for all cases but *φ* = *b*, were fitted to a normal distribution. For *φ* = *b*, the fitted values formed a long-tailed distribution (Figure 5) and were accordingly fitted to a log-normal distribution. Fitting was performed by assuming that fitted *φ*_*i*_ values are samples drawn from a normal distribution with mean *φ*_0_ and standard variation *σ*_*φ*_ (for *φ* = *b*, it was assumed that ln(*φ*) is drawn from a normal distribution)

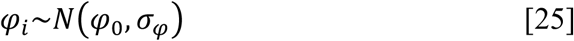

where *φ*_0_ and *σ*_*φ*_ were optimized by maximizing

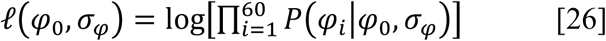

and *P*(*φ*_*i*_*φ*_0_, *σ*_*φ*_) was calculated using a normal distribution. To assess the fit quality, normality test was performed for (*φ*_*i*_ − *φ*_0_)/*σ*_*φ*_ using the Kolmogorov-Smirnov test against standard normal distribution, with mean 0 and standard deviation 1 (the log of fitted values were tested for normality for *φ* = *b*). For all cases, the null hypothesis that the optimized *φ*_*i*_ values are drawn from a normal distribution could not be rejected. The fitted normal distributions are shown in Figures 4 and 5, and the normal distributions parameters and Kolmogorov-Smirnov test *p* -values are given in Tables 1 and 2.

**Figure 4.**
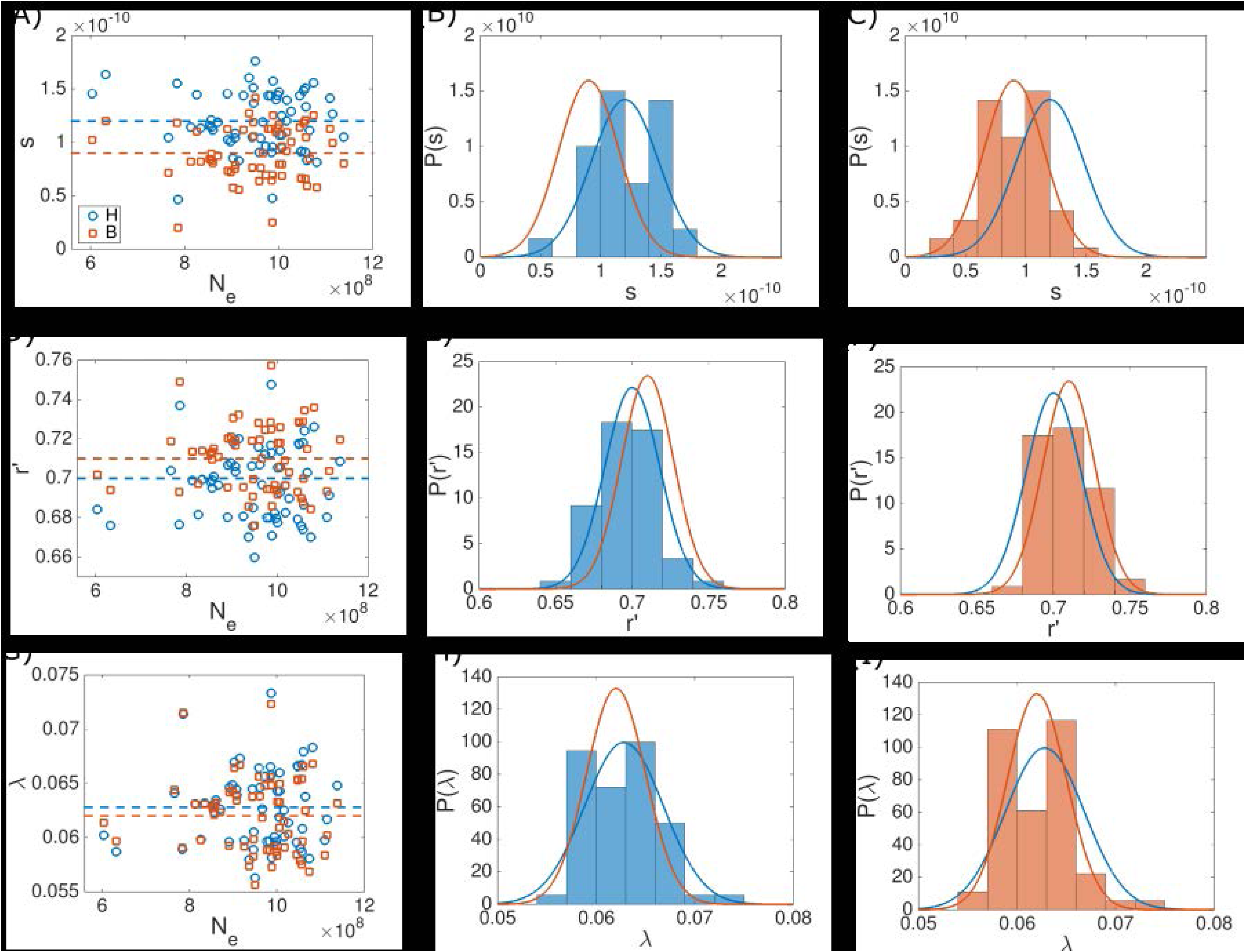
Fitted latent variable values under the power law deletion bias model (Eq.(14)) for *φ* = *s* (A-C), *φ* = *r*′ (D-F) and *φ* =*λ* (G-I). The fits were obtained using the hard fitting methodology (blue) and the hierarchical Bayesian model (orange). Fitted *φ* values for all ATGCs are plotted against the effective population size in the leftmost column. The mean values of the distributions are indicated by dashed lines. The fitted *φ* values histograms are shown together with the latent variable distributions, which are indicated by solid lines. The distribution parameters are given in Table 1. Histograms obtained using the hard fitting methodology are shown in the middle column, and histograms obtained under the hierarchical Bayesian model are shown in the rightmost column.

**Figure 5.**
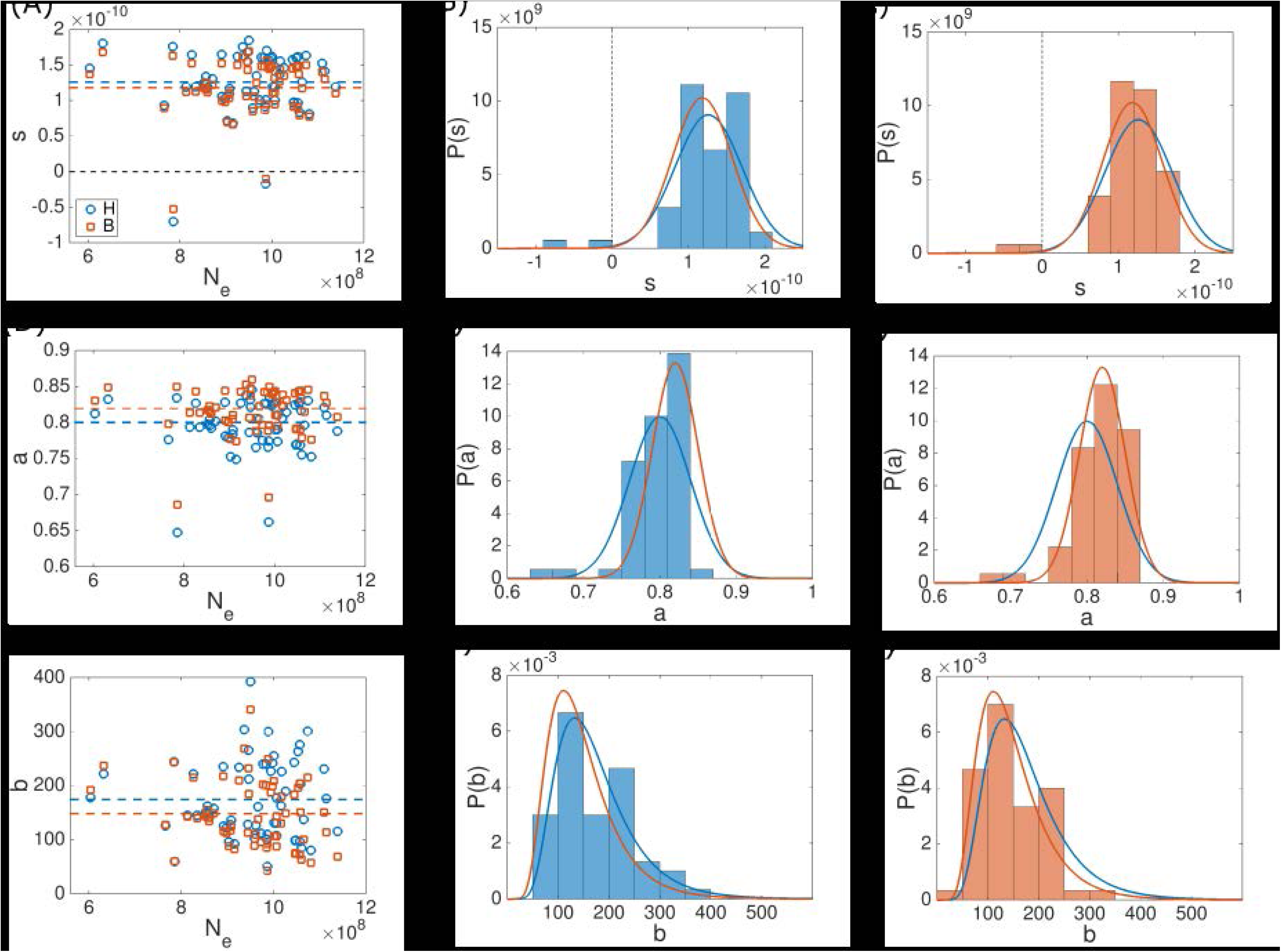
Fitted latent variable values under the linear deletion bias model (Eq.(17)) for *φ* = *s* (A-C), *φ* = *a* (D-F) and *φ* = *b* (G-I) The fits were obtained using the hard fitting methodology (blue) and the hierarchical Bayesian model (orange). The fitted *φ* values for all ATGCs are plotted against the effective population size in the leftmost column. Values are indicated by markers and mean values of the distributions are indicated by dashed lines. Fitted *φ* values histograms are shown together with latent variable distributions, which are indicated by solid lines. The parameters of the distributions are given in Table 2. Histograms obtained using the hard fitting methodology are shown in the middle column, and histograms obtained using the hierarchical Bayesian model are shown in the rightmost column.

In the ad-hoc hard fitting method described above, Eq.(7) was used to adjust latent variable values such that the model distributions centered around the observed genome sizes. The fitted latent variable values are then scattered around some typical value (Figures 4 and 5).

Moreover, fitted values form distributions that are statistically indistinguishable from normal distributions (with the exception of the case *φ* = *b*, which forms a log-normal distribution). It is possible to rely on this observation and implement an alternative optimization methodology, where a prior distribution *P*_*φ*_ is assumed for the latent variable. In the following, normal distributions were assumed as priors, with the exception of a log-normal distribution for the case when *b* is set as the latent variable. Accordingly, a specific value *φ*_*i*_ of the latent variable is associated with a probability *P*_*φ*_(*φ*_*i*_|*φ*_0_, *σ*_*φ*)_. The probability of the observed genome sizes *x*_*ii*_ for species *j* of ATGC *i* can be then calculated using the Bayes rule, and is given by

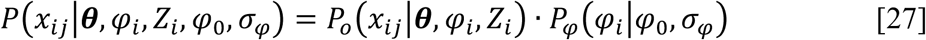

This formulation is known as the hierarchical Bayesian model (Gelman et al., 1995). The probability of *x*_*ij*_ depends on the prior distribution of *φ*_*i*_ parameters (*φ*_0_ and *σ*_*φ*_) indirectly, and in an hierarchical manner: *x*_*ii*_ depends directly on *φ*_*i*_, which in turn occurs with the probability *P*_*φ*_ that depends on *φ*_0_ and *σ*_*φ*_. The prior distribution parameters are optimized as well during the fitting process and are therefore included in the set of global parameters ***θ***. The log-likelihood is then given by ℓ(***θ***, *φ*)

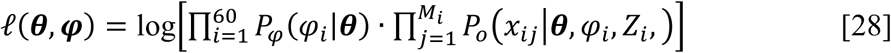

where *x*_*ij*_ is observed genome size for species *j* out of *M*_*i*_ species of ATGC *i*. In more compact way, the equation above can be written as

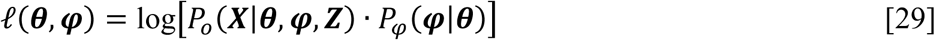

Note that within this formulation, the maximization of ℓ(***θ***, ***φ***) is performed in a 64-dimensional parameter space (60 ***φ*** latent variable values, 2 global model parameters ***θ*** and 2 parameters describing the prior distribution *P*_*φ*_ of the latent variable). However, for the optimization of ***θ***, it is possible to sum over all possible values of the latent variable ***φ***, such that *P*_***θ***_(***X***|***θ***, ***Z***) of Eq.(1) is given by

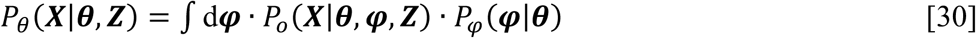

and the optimization of ***θ*** is performed by maximizing ℓ(***θ***). To test the validity of the hierarchical Bayesian approach, when applied using the population-genetic model of genome evolution, 9 realizations of artificial ATGCs ware generated using the distribution of genome sizes given by the model (Eq.(7); see Methods for details). The realizations were generated using parameter values similar to the fitted parameters obtained using the hard fitting methodology. We then applied the hierarchical Bayesian model fitting algorithm to the artificial ATGCs and verified that the optimized parameters values were similar to those of the parameters used for generating the artificial ATGCs (Figure S2). In all realizations, the *λ* value was inferred to a good accuracy, with a tendency for the fitting values to be slightly lower than the actual ones. The fitted values of *s* and *r*′ typically have larger errors because variation of *s* can be compensated by the variation of *r*′, and vice versa. Accordingly, the fitted values of *s* and *r*′ follow a line (Figure S2D). However, the under-estimation of *λ* is compensated by slightly greater values of *r*′, resulting in a slight offset of the *s* − *r*′ trend line with respect to the actual values. Finally, the hierarchical Bayesian model was applied to optimize model parameters according to the genomic data, where one genome size model parameter is set as latent variable. Fitted values of global parameters ***θ*** are summarized in Table 1 and Table 2, where global parameters now include the parameters of the prior distribution of the latent variable, *φ*_0_ and *σ*_*φ*_. Using these optimized ***θ*** values together with Eq.(28) allows fitting the ATGC-specific *φ* values (Figure 4 and Figure 5). As with the ad-hoc hard-fitting methodology, there was no significant correlation between fitted ***φ*** values and *N*_*e*_ (see Table 1 and Table 2), with the exception of *φ* = *b*, where the Spearman’s correlation coefficient is ρ = -0.24 with *p* -value 0.06. Notably, both optimization methodologies gave similar results in terms of the optimized values of ***θ*** and *φ*, as shown in Table 1, Table 2 and Figures 4 and 5.

In all cases, the genome size distributions produced by the model centered on the observed genome sizes, either by design, as in the hard-fitting algorithm, or as a result of maximizing the log-likelihood, as in the hierarchical Bayesian approach. However, the observed widths of the genome size distributions are not predicted perfectly by the model, as shown in Figure 3. It is therefore natural to consider the case where more than one model parameter is set as a latent variable. Although generalizing the hierarchical Bayesian model to account for more than one latent variable is straightforward, the calculation of the integral of Eq.(30) is computationally intensive for more than one latent variable. However, it is possible to explore a setting with more than one latent variable in the hierarchical Bayesian model that is expected to produce similar results. As the calculation of the integral in Eq.(30) requires long computation times, the assessment is performed using the expression for ℓ(***θ***, ***φ***) of Eq.(28). Specifically, for deletion bias modelled as in Eq.(14), all three genome size model parameters (*s*, *λ* and *r*′) are set as latent variables, and the normal distributions fitted to the latent variables values obtained by applying the hard-fitting methodology are used as priors. Prior distributions are not optimized such that the product term of Eq.(28) can be calculated separately for each ATGC, with high efficiency. It is important to note that this is an approximation because the prior distributions that are used here were obtained when optimizing one latent variable at a time. Another possibility is to perform the optimization in the 64 dimensional parameter space of ℓ(***θ***, ***φ***) in two stages: for the given ***θ*** values, latent variables are fitted for each ATGC separately such that ℓ(***θ***, *φ*_*i*_) is maximized. This approach was applied for *φ* = {*λ*, *r*′}. Both assessments produced results similar to those obtained for one latent variable, so we conclude that, within the current modeling framework, the agreement between the model and the observed genome size distributions cannot be significantly improved further by considering additional latent variables under the hierarchical Bayesian model.

Finally, the distributions for the latent variable can be used in order to derive estimations for maximum and minimum genome sizes. The optimized ***θ*** values together with *φ* values from the optimized prior distributions tails were substituted into the model approximation for mean genome size of Eq.(8). Specifically, *φ* values from percentiles 1 to 10 and 90 to 99 were used, where each of the two ranges corresponds either to the maximum or to the minimum genome size estimates, depending on the choice of the latent variable. For example, when *φ* = *λ* or *φ* = *r*′, the left tail of the distribution (1 to 10 percentile) corresponds to the maximum genome size estimates, whereas for all other choices of *φ*, the left tail corresponds to the minimum genome size estimates. The effective population size was set arbitrarily to *N*_*e*_ = 10^9^. Estimations for *φ* = *s*, *φ* = *λ*, *φ* = *r*′ and *φ* = *a* are shown in Figure 6. For deletion bias modeled by Eq.(14), the estimates are roughly consistent with the observed minimum and maximum genome sizes of prokaryotes (excluding the smallest genomes of intracellular parasitic bacteria) (Koonin and Wolf, 2008). Notably, genome size diverges for the deletion bias of Eq.(17) with *φ* = *s* or *φ* = *a* as a latent variable. The deletion bias of Eq.(17) results from linear approximations for the acquisition and the deletion rates. Accordingly, gain and loss rates are linear with respect to genome size, where the slope of *P*_+_ is smaller than the slope of *P*_-_, albeit with a non-zero intercept (model parameter *b*). A finite genome size *x*_0_, where *P*_+_ = *P*_-_ therefore exists, and the condition of Eq.(10) is satisfied. However, for *a* = *e*^−*N*_*e*_*s*^, both rates, *P*_+_ and *P*_-_, have the same slope and *P*_+_ > *P*_-_ for all genome sizes, such that the genome size diverges.

**Figure 6.**
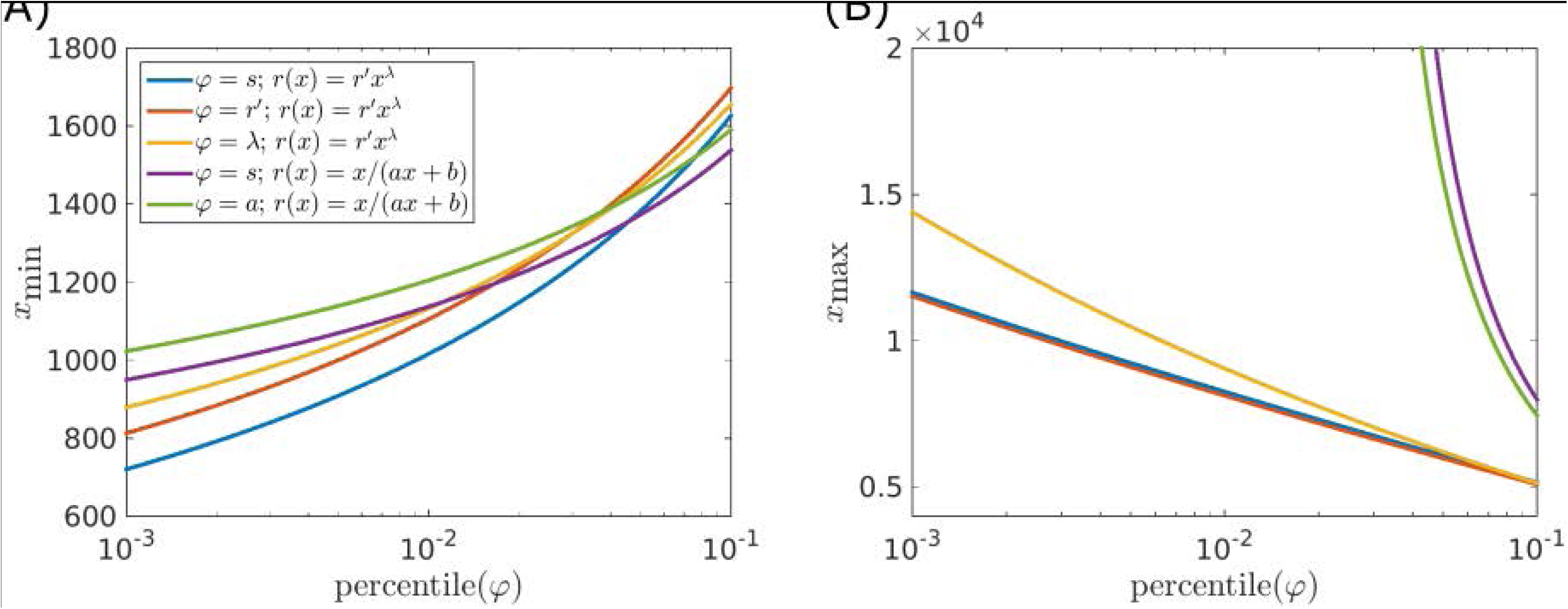
Maximum and minimum equilibrium genome sizes calculated using Eq.(8) with parameters fitted under the hierarchical Bayesian model. Latent variables and deletion bias models are indicated in the inset. The effective population size was set as *N*_*e*_ = 10^9^. For each fit, the latent variable was taken from the left tail (percentiles 1-10) or the right tail (percentiles 90-99) of the optimized distribution of the latent variable. All estimates for maximum or minimum genome sizes, based on the different choices of the latent variable, are plotted together. As a result the same figure mixes distributions left and right tail for different choices of *φ*. (A) For *φ* = *r*′ and *φ* = *λ* the *x* axis indicates 1 − *P*, where *P* is the percentile. (B) For *φ* = *s* and *φ* = *a* the *x* axis indicates 1 − *P*, where *P* is the percentile.

## Discussion

Our previous effort on modeling microbial genome evolution (Sela et al., 2016) has shown that for all ATGCs, the best fit between the model-generated and observed distributions of genome sizes were obtained with positive *s* values and r> 1 (deletion bias). Given that the deletion bias indeed appears to be a universal characteristic of genome evolution (Kuo and Ochman, 2009; Petrov, 2002; Petrov et al., 2000), we have concluded that prokaryotic genomes typically evolve under a selection-mutation balance regime as opposed to a streamlining regime. In biologically oriented terms, these results seem to indicate that, on average, benefits of new genes acquired by microbial genomes outweigh the cost of gene maintenance and expression. However, the actual values of the selection coefficient yielded by the model were extremely low, on the order of 10^-12^, suggesting that the selection affecting an average gene was extremely weak, but also that these values could be under-estimates. The latter possibility was additionally suggested by the observation that, although the model yielded good fits for the means of the genome size distributions, the width of the distributions was significantly over-estimated (Figure 3A). In the previous study, we made the strong assumption that the parameters of microbial genome evolution were universal across the entire prokaryotic diversity represented in the ATGCs. The results indicate that the contribution of the universal factors is indeed substantial but fails to account for all or even most of the variation in the genome size distributions indicating that, not unexpectedly, ATGC-specific factors are important for genome evolution as well.

In the present work, we attempted to take into account the group-specific evolutionary factors by using two independent optimization approaches. Both procedures were used together with two different functional forms of the deletion bias. In all cases, the results were similar, with *s*∼10^− 10^, *λ*∼0.06 and *r*′∼0.7 for a power law deletion bias (Table 1), and *s*∼10^− 10^, *a*∼0.8 and *b*∼175 for a deletion bias based on linear acquisition and deletion rates (Table 2). Introducing latent variables allowed incorporation of ATGC-specific effects into the fitting process. However, variation in one model parameter can be compensated by adjustment of another model parameter, such that all fits are similar in terms of log-likelihood and thus it is impossible to disambiguate global from local factors affecting the evolution of genome size in terms of model parameters. Nevertheless, the optimized values of the latent variables form relatively narrow distributions around the means (Figures 4 and 5), such that, for the deletion bias of Eq.(14), the ratios between standard deviation and mean values are 0.28, 0.06 and 0.03 for *φ* = *s*, *φ* = *λ* and *φ* = *r*′, respectively. For the linear deletion bias given by Eq.(17), the ratios between standard deviation and mean values are 0.35, 0.05 and 0.46 for *φ* = *s*, *φ* = *a* and *φ* = *b*, respectively. In both cases, the higher value among those obtained with the hard fitting and the hierarchical Bayesian model methodologies is indicated. Thus, the mean values give good estimates for model parameters for all ATGCs. The mean selection coefficient of *s*∼10^− 10^ associated with the gain of one gene implies that, on average, acquisition of a gene is beneficial, and that microbial genomes typically evolve under a weak selection regime, with the characteristic selection strength *N*_*e*_ *s*∼0.1. In highly abundant organisms, transition to a strong selection regime, with *N*_*e*_ *s* > 1.0, appears possible. These values of *s* appear to be substantially more realistic than the lower values obtained previously, indicating that global and group-specific evolutionary factors synergistically affect microbial genome evolution. This result is consistent with the observed significant, positive correlation between the genome size and selection strength on the protein level and appears intuitive given the diversity of bacterial lifestyles that conceivably drives adaptive gene acquisition. The selective pressure towards larger genomes, manifested in the positive selection coefficients, is balanced by the deletion bias, which is consistently greater than unity. Notably, an independent duplication-loss-transfer model of microbial evolution that we have developed recently in order to compare the evolutionary regimes of different classes of genes has yielded closely similar mean values of the selection coefficient (Iranzo et al., 2017).

In this work, the deletion bias is considered genome size-dependent and is modelled as a power law or as the ratio of linear approximations for the acquisition and the deletion rates. We found that the best fitted power value is *λ*∼0.06. This value indicates that the genome size dependencies of gene acquisition and deletion rates are generally similar but the deletion rate grows slightly faster with the genome size. This difference, although slight, could put a limit on microbial genome growth. Estimates for minimal and maximal genome sizes were derived using model parameters from the edges of latent variables distributions (percentiles 1% and 99%). The estimations derived using a power law deletion bias were consistent with the observed prokaryotic genome sizes, genome size diverged when considering values from the edges of the distributions together with a linear approximation for the deletion bias. This divergence suggests that the linear approximation for the acquisition and deletion rates holds only locally, and breaks down when a wide range of parameters is considered.

Given the compensation between the *s* and *r’* values, the comparison between the values of these parameters obtained for different ATGCs should be approached with caution. Nevertheless, with this caveat, it is worth noting that the lowest mean values of the selections coefficient were estimated for parasitic bacteria with degraded genomes, such as *Mycoplasma* and *Chlamydia*, whereas the highest values were obtained for complex environmental bacteria with large genomes, such as *Rhizobium* and *Serratia* (Supplementary Tables 2 and 3). These differences are compatible with the proposed regime of adaptive evolution of microbial genomes under (generally) weak selection for functional diversification.

## References

Doolittle, W. F. 1999. Lateral genomics. Trends Cell Biol 9: M5–8.

Edgar, R. C. 2004. MUSCLE: multiple sequence alignment with high accuracy and high throughput. Nucleic Acids Res 32: 1792–1797.

Gelman, A., J. Carlin, H. Stern and D. Rubin 1995. Bayesian Data Analysis. New York: Chapman and Hall.

Hurst, L. D. 2002. The Ka/Ks ratio: diagnosing the form of sequence evolution. Trends Genet 18: 486.

Iranzo, J., J. A. Cuesta, S. Manrubia, M. I. Katsnelson and E. V. Koonin 2017. Disentangling the effects of selection and loss bias on gene dynamics. Proc Natl Acad Sci U S A in press.

Koonin, E. V. 2003. Comparative genomics, minimal gene-sets and the last universal common ancestor. Nature Rev. Microbiol. 1: 127–136.

Koonin, E. V. 2009. Evolution of genome architecture. Int J Biochem Cell Biol 41: 298–306.

Koonin, E. V., K. S. Makarova and L. Aravind 2001. Horizontal gene transfer in prokaryotes: quantification and classification. Annu Rev Microbiol 55: 709–742.

Koonin, E. V. and Y. I. Wolf 2008. Genomics of bacteria and archaea: the emerging dynamic view of the prokaryotic world. Nucleic Acids Res 36: 6688-6719. doi: gkn668 [pii] 10.1093/nar/gkn668

Kristensen, D. M., Y. I. Wolf and E. V. Koonin 2017. ATGC database and ATGC-COGs: an updated resource for micro- and macro-evolutionary studies of prokaryotic genomes and protein family annotation. Nucleic Acids Res 45: D210-D218. doi: 10.1093/nar/gkw934 gkw934 [pii]

Kryazhimskiy, S. and J. B. Plotkin 2008. The population genetics of dN/dS. PLoS Genet 4: e1000304. doi: 10.1371/journal.pgen.1000304

Kuo, C. H., N. A. Moran and H. Ochman 2009. The consequences of genetic drift for bacterial genome complexity. Genome Res 19: 1450-1454. doi: 10.1101/gr.091785.109 gr.091785.109 [pii]

Kuo, C. H. and H. Ochman 2009. Deletional bias across the three domains of life. Genome Biol Evol 1: 145-152. doi: 10.1093/gbe/evp016

Lynch, M. 2007. The origins of genome archiecture. Sunderland, MA: Sinauer Associates.

Lynch, M. 2006. Streamlining and simplification of microbial genome architecture. Annu Rev Microbiol 60: 327–349.

Lynch, M. and J. S. Conery 2003. The origins of genome complexity. Science 302: 1401–1404.

Lynch, M. and G. K. Marinov 2015. The bioenergetic costs of a gene. Proc Natl Acad Sci U S A 112: 15690-15695. doi: 10.1073/pnas.1514974112 1514974112 [pii]

McCandlish, D. M., C. L. Epstein and J. B. Plotkin 2015. Formal properties of the probability of fixation: identities, inequalities and approximations. Theor Popul Biol 99: 98-113. doi: 10.1016/j.tpb.2014.11.004 S0040-5809(14)00094-X [pii]

Mira, A., H. Ochman and N. A. Moran 2001. Deletional bias and the evolution of bacterial genomes. Trends Genet 17: 589–596.

Novichkov, P. S., I. Ratnere, Y. I. Wolf, E. V. Koonin and I. Dubchak 2009a. ATGC: a database of orthologous genes from closely related prokaryotic genomes and a research platform for microevolution of prokaryotes. Nucleic Acids Res 37: D448-454. doi: 10.1093/nar/gkn684 gkn684 [pii]

Novichkov, P. S., Y. I. Wolf, I. Dubchak and E. V. Koonin 2009b. Trends in prokaryotic evolution revealed by comparison of closely related bacterial and archaeal genomes. J Bacteriol 191: 65–73.

Pal, C., B. Papp and M. J. Lercher 2005. Adaptive evolution of bacterial metabolic networks by horizontal gene transfer. Nat Genet 37: 1372-1375. doi: ng1686 [pii] 10.1038/ng1686

Petrov, D. A. 2002. DNA loss and evolution of genome size in Drosophila. Genetica 115: 81–91.

Petrov, D. A., T. A. Sangster, J. S. Johnston, D. L. Hartl and K. L. Shaw 2000. Evidence for DNA loss as a determinant of genome size. Science 287: 1060-1062. doi: 8235 [pii]

Puigbo, P., A. E. Lobkovsky, D. M. Kristensen, Y. I. Wolf and E. V. Koonin 2014. Genomes in turmoil: quantification of genome dynamics in prokaryote supergenomes. BMC Biol 12: 66. doi: 10.1186/S12915-014-0066-4 S12915-014-0066-4 [pii]

Sela, I., Y. I. Wolf and E. V. Koonin 2016. Theory of prokaryotic genome evolution. Proc Natl Acad Sci U S A 113: 11399-11407. doi: 1614083113 [pii] 10.1073/pnas.1614083113

Treangen, T. J. and E. P. Rocha 2011. Horizontal transfer, not duplication, drives the expansion of protein families in prokaryotes. PLoS Genet 7: e1001284. doi: 10.1371/journal.pgen.1001284

Yang, Z. 2007. PAML 4: phylogenetic analysis by maximum likelihood. Mol Biol Evol 24: 1586–1591.

